# Molecular evolution of non-visual opsin genes across environmental, developmental, and morphological adaptations in frogs

**DOI:** 10.1101/2022.11.10.515783

**Authors:** John L Boyette, Rayna C Bell, Matthew K Fujita, Kate N Thomas, Jeffrey W Streicher, David J Gower, Ryan K Schott

**Affiliations:** Department of Biological Sciences, The University of Alabama, Tuscaloosa, Alabama, USA; Department of Biology, Berry College, Rome, Georgia, USA; Department of Vertebrate Zoology, National Museum of Natural History, Smithsonian Institution, Washington DC, USA; Department of Herpetology, California Academy of Sciences, San Francisco, California, USA; Systematics and Biodiversity Science Cluster, National Science Foundation, Alexandria, Virginia, USA; Department of Life Sciences, Natural History Museum, London, UK; Department of Biology, York University, Toronto, Ontario, CAN

**Keywords:** Amphibia, Anura, eyes, selection, transcriptome, visual ecology

## Abstract

Non-visual opsins are transmembrane proteins expressed in the eyes, skin, and brain of many animals. When paired with a light-sensitive chromophore, non-visual opsins form photopigment systems involved in various non-visual, light-detection functions, including circadian rhythm regulation, light-seeking behavior, and detection of seasonality. Previous research has primarily explored the diversity and function of non-visual opsins in model organisms, with few studies investigating their molecular evolution in non-model species. Here we explored molecular evolution of non-visual opsin genes in anuran amphibians (frogs and toads). With diverse lifestyles ranging from fossorial to aquatic, anurans inhabit a diverse array of light environments, which makes them a compelling system for studying the evolution of light detection mechanisms. Using whole-eye transcriptomes from 79 anuran species, as well as genomes and multi-tissue transcriptomes from an additional 15 species, we 1) identify which non-visual opsin genes are expressed in the eyes of anurans; 2) compare selective constraint (ω, or dN/dS) among non-visual opsin genes; and 3) test for potential adaptive evolution by comparing selection between discrete ecological classes in anurans (e.g. diurnal versus non-diurnal). We consistently recovered 14 non-visual opsin genes from anuran eye transcriptomes, compared to 18 genes that we recovered genome wide, and detected positive selection in a subset of these genes. We also found variation in selective constraint between discrete ecological and life-history classes, which may reflect functional adaptation in non-visual opsin genes. Although non-visual opsins remain poorly understood, these findings provide insight into their molecular evolution and set the stage for future research on their potential function across taxa with diverse environmental, developmental, and morphological adaptations.

## 1 Introduction

Animals rely on light detection to accomplish many biologically critical functions. Visual photosensitivity allows animals to acquire food, locate mates, and avoid predators. In recent years, the role of non-visual photosensitivity has received increasing attention, revealing a diverse suite of physiologically important non-visual functions including calibration of circadian rhythm, regulation of light-seeking behavior, and initiation of seasonal reproductive changes (Göz et al., 2008; Fernandes et al., 2012; Nakane et al., 2010; Nakane, Shimmura, Abe, & Yoshimura, 2014). The basis of animal photosensitivity lies in the conversion of light stimuli into neural stimuli – a process known as phototransduction – which is initiated by photopigments embedded in the membrane of light sensitive cells. Each photopigment is composed of a transmembrane opsin protein encapsulating a light sensitive chromophore. Different opsins confer distinct spectral sensitivities to their respective photopigments by maximally absorbing a specific wavelength of light. Upon absorption of light, the chromophore photoisomerizes, undergoing a conformational change that initiates phototransduction, generating a neural signal that can be interpreted for either visual or non-visual functions (Terakita, 2005; Hunt & Collin, 2014). The function of opsins in initiating phototransduction is illustrated in Figure 1.

**FIGURE 1.**
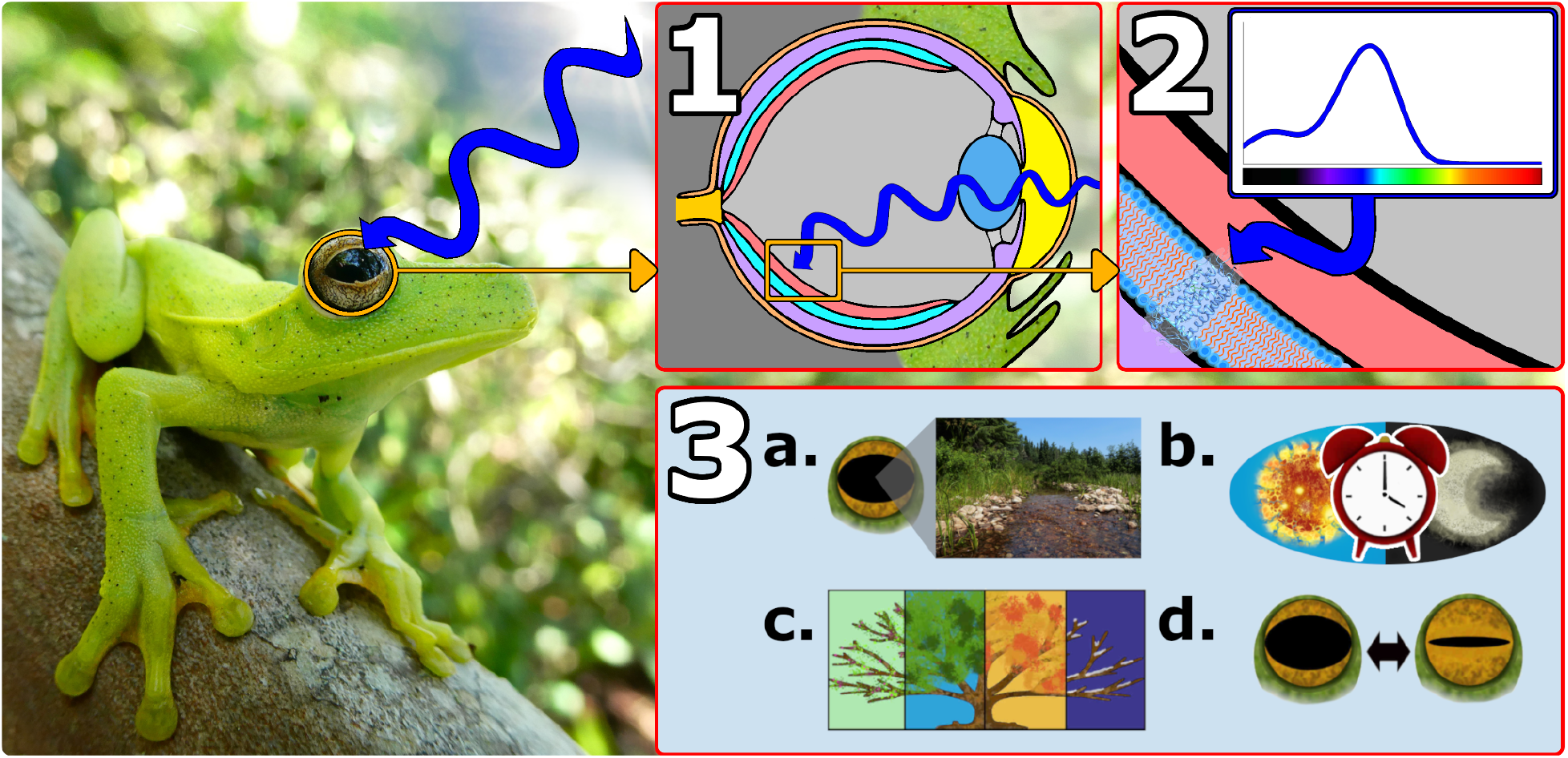
General overview of opsin function in the eye of a frog (*Hypsiboas albomarginatus* pictured here). **1)** Light enters the eye and is focused on the retina. **2)** Light reaches a photopigment embedded in the membrane of a light-sensitive retinal cell. The opsin maximally absorbs a specific wavelength of light. In this example, the opsin maximally absorbs blue light. Absorption of light stimulates photoisomerization of the chromophore encapsulated within the opsin. **3)** A neural signal is generated and sent to the brain to be interpreted for visual or non-visual purposes. These may include (**a.**) formation of a visual image, (**b.**) calibration of circadian rhythm, (**c.**) detection of seasonality, or (**d.**) regulation of the pupillary light response.

Opsins are divided broadly into eight groups based on amino-acid sequence similarity, molecular function, and signaling properties. The vertebrate visual opsin group contains opsins involved in initiating the formation of visual images, including rod opsins (RH1 and RH2) and cone opsins (LWS, SWS1, and SWS2). These opsins are associated with “bleaching” photopigments, meaning that following light exposure and photoisomerization, the chromophore dissociates from the opsin and renders the photopigment unreactive until the chromophore can be regenerated (Tsukamoto, 2014). Opsins in the retinal photoisemrase group (RGR) and the peropsin group (RRH) function to regenerate chromophores used by bleaching photopigments (Terakita, 2005; Radu et al., 2008; Zhang et al., 2019). The other opsin groups include the vertebrate non-visual opsin group (PAR, PARA, VAOP, and PIN), the encephalopsin/tmt-opsin group (OPN3 and TMT-), the Gq-coupled opsin/melanopsin group (OPN4-), the Go-coupled opsin group, and the neuropsin group (NEUR-). These groups are generally found in non-visual photopigments expressed in various major organs including the eye, brain, and skin of many organisms (e.g., Davies et al., 2015; Foster & Bellingham, 2004; Kelley & Davies, 2016; Nakane et al., 2010). These opsin groups function in either bleaching or “non-bleaching” photopigments. Non-bleaching photopigments, also referred to as “bistable” photopigments, retain their chromophore following light exposure and photoisomerization. This is relevant to non-visual photopigments because many are expressed in extraocular tissues that lack the specialized chromophore-regeneration mechanisms of the retina (Tsukamoto, 2014; Steindal & Whitmore, 2020). For the purposes of this study, we broadly refer to all opsins outside the vertebrate visual opsin group as non-visual opsins.

The scope of opsin diversity coupled with the breadth of tissues and taxa expressing these genes suggests that the biological relevance of light detection extends far beyond the visual system. Among vertebrates, teleost fishes demonstrate the greatest opsin diversity, with 10 visual and 32 non-visual opsins reported in zebrafish (Davies et al., 2015). This diversity likely arose through whole-genome duplication events, and the retention of these opsin genes in zebrafish is hypothesized to confer an adaptive advantage in dynamic freshwater light environments. Mammals, on the other hand, have lost multiple opsins, with only three visual and six non-visual opsins reported in mice (Davies et al., 2015; Gemmell et al., 2020). This disparity of opsin diversity across vertebrates emphasizes the importance of ecology in the evolution of opsin systems. For example, the loss of opsin diversity in mammals is hypothesized to result from a “nocturnal bottleneck” in which ancestral mammals transitioned to nocturnal lifestyles and encountered a reduced need for broad spectral sensitivity (Gerkema, Davies, Foster, Menaker, & Hut, 2013; Borges et al., 2018). Ecological transitions to low-light environments are also used to explain loss of opsin diversity in many other amniote taxa, including geckos (Pinto, Nielsen, & Gamble, 2019), crocodilians (Emerling, 2017), snakes (Davies et al., 2009; Gower, Hauzman, Simões, & Schott, 2022; Schott, Van Nynatten, Card, Castoe, & Chang, 2018), whales (Meredith, Gatesy, Emerling, York, & Springer, 2013), burrowing rodents (Emerling & Springer, 2014), and nocturnal primates (Kawamura & Kubotera, 2004). These examples highlight the impact of low-light ecological transitions on opsin diversity and evolution; however, the influence of other environmental, developmental, and morphological adaptations remains poorly studied.

Frogs and toads (Anura, hereafter collectively “frogs”) provide an opportune system in which to investigate opsin diversity and evolution because they demonstrate remarkable variation in activity period, habitat, distribution, life history, and pupil shape (e.g. Moen, Irschick, & Wiens, 2013; Thomas et al., 2021; Wiens, Graham, Moen, Smith, & Reeder, 2006). This variation exposes frogs to diverse light environments and sensory constraints, which in turn introduce unique evolutionary challenges to the non-visual system. For example, the nocturnal bottleneck hypothesis exemplifies a connection between evolution of non-visual opsins and adaptation to new activity periods and habitats. We also expect that species distribution has influenced non-visual opsin evolution, because species distributed outside tropical zones experience more predictable seasonal variation in photoperiod (Borah, Renthlei, & Trivedi, 2019; Canavero & Arim, 2009). Variation in seasonality and photoperiod both have implications for non-visual opsin evolution because these proteins are involved in responses to seasonality (Nakane et al., 2010) and regulation of circadian rhythm (Göz et al., 2008). Many frog species also experience a dramatic shift in light environment across development as they metamorphose from aquatic larvae to terrestrial adults. This biphasic life history potentially subjects non-visual opsins to disparate environmental constraints and selective pressures across metamorphosis, resulting in adaptive decoupling in larval developing frog species (Schott et al., 2021). However, a subset of frog species, known as direct-developers, lack a free-living aquatic larval stage. Direct-developers provide an opportunity to investigate the role of non-visual opsins across life-history strategies, which is worthwhile because these proteins are involved in regulating light-seeking behavior in aquatic larvae (Fernandes et al., 2012). Finally, frogs demonstrate a strikingly diverse suite of pupil shapes, which each regulate the amount of light reaching the retina through pupillary constriction (Malmström & Kröger, 2006; Thomas et al., 2021). This morphological diversity may have implications for non-visual opsin evolution because these proteins have been implicated in regulation of pupillary light responses (Keenan et al., 2016). Taken together, the environmental, developmental, and morphological diversity of frogs makes them an attractive study system in which to investigate non-visual opsin diversity and evolution.

Here, we extract non-visual opsin genes from *de novo* whole-eye transcriptome assemblies of 79 frog species, as well as genomes and multi-tissue transcriptomes of an additional 15 species. Together, these 94 frog species represent 34 of 56 currently recognized families, including a broad sampling of environmental, developmental, and morphological adaptations, and we predict that this variation has influenced the diversity and molecular evolution of the non-visual opsins. We aim to 1) identify which non-visual opsin genes are expressed in the eyes of frogs; 2) compare selection among non-visual opsin genes; and 3) test for potential adaptive evolution by comparing selection among frogs with differing ecologies.

## 2 Materials and Methods

### 2.1 Species sampling

Our sampling included a total of 94 species, including 88 whole eye transcriptomes from 79 species and 19 genomes or multi-tissue transcriptomes from 15 additional species. Of the 88 whole eye transcriptomes, 79 of these samples were collected from adult frogs, and the remaining nine were collected from larval frogs (Supplementary Table S2). Frogs were sampled from wild populations in Australia, Brazil, Cameroon, Ecuador, Equatorial Guinea, French Guiana, Gabon, the Seychelles, the United Kingdom, and the United States (Supplementary Table S2). Additional species were obtained from commercial dealers or captive colonies. Most individuals were kept in complete darkness (i.e., were dark adapted) for 3+ hours prior to euthanasia (via immersion in a solution of MS222) because one eye was removed for microspectrophotometry measurements as part of another study (Thomas et al. in prep). Whole eyes were extracted, punctured and placed in RNAlater (Ambion) for at least 24 hours at 4°C to allow the RNALater to saturate the cells, prior to freezing and storage at −80°C until use. Some samples were collected at remote field sites and were kept as cool as possible in RNAlater prior to freezing at −80°C at the earliest opportunity. Voucher specimens and tissues for further genetic analysis were accessioned in natural history museums (Supplementary Table S2). To supplement the phylogenetic and ecological diversity of species we collected, we searched Genbank for publicly available frog genomes and multi-tissue transcriptomes to include in our analyses. These included 19 genomes or multi-tissue transcriptomes from adult, larval, or mixed adult/larval frog samples (Supplementary Table S3). Our combined sampling includes 94 species, representing 34 of 56 currently recognized frog families (Frost, 2021; Figure 3).

**FIGURE 2.**
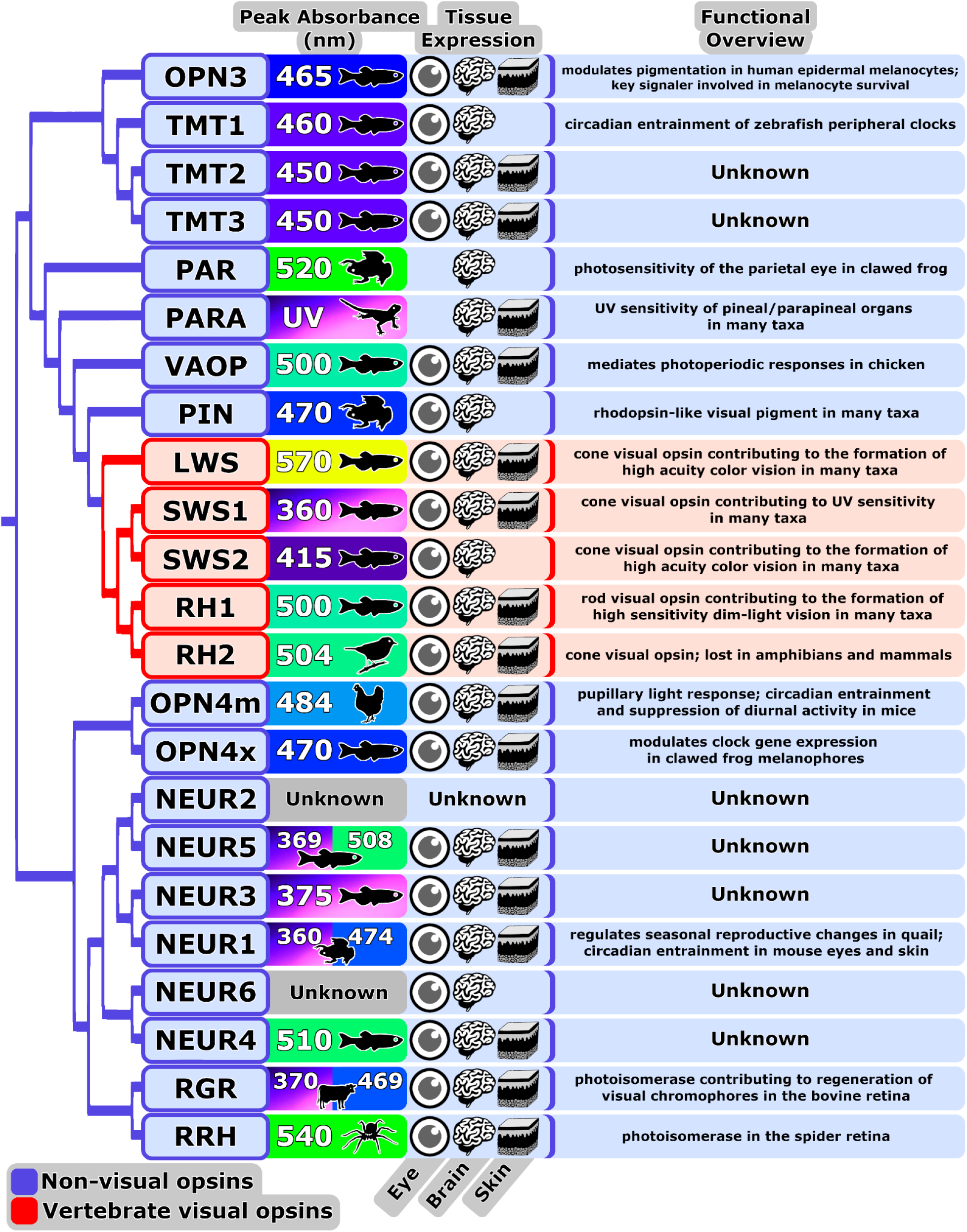
Diversity of spectral sensitivity, tissue expression, and function across the visual and non-visual opsins. Phylogenetic hypothesis based on Beaudry et al., (2017) and Davies et al., (2015). Peak absorbance measurements and corresponding color are displayed beside the taxa in which each measurement was observed. Tissue expression based on transcriptomic profiles in zebrafish (Davies et al., 2015). Citations for peak absorbance and functional overview notes can be found in Supplementary Table S1.

**FIGURE 3.**
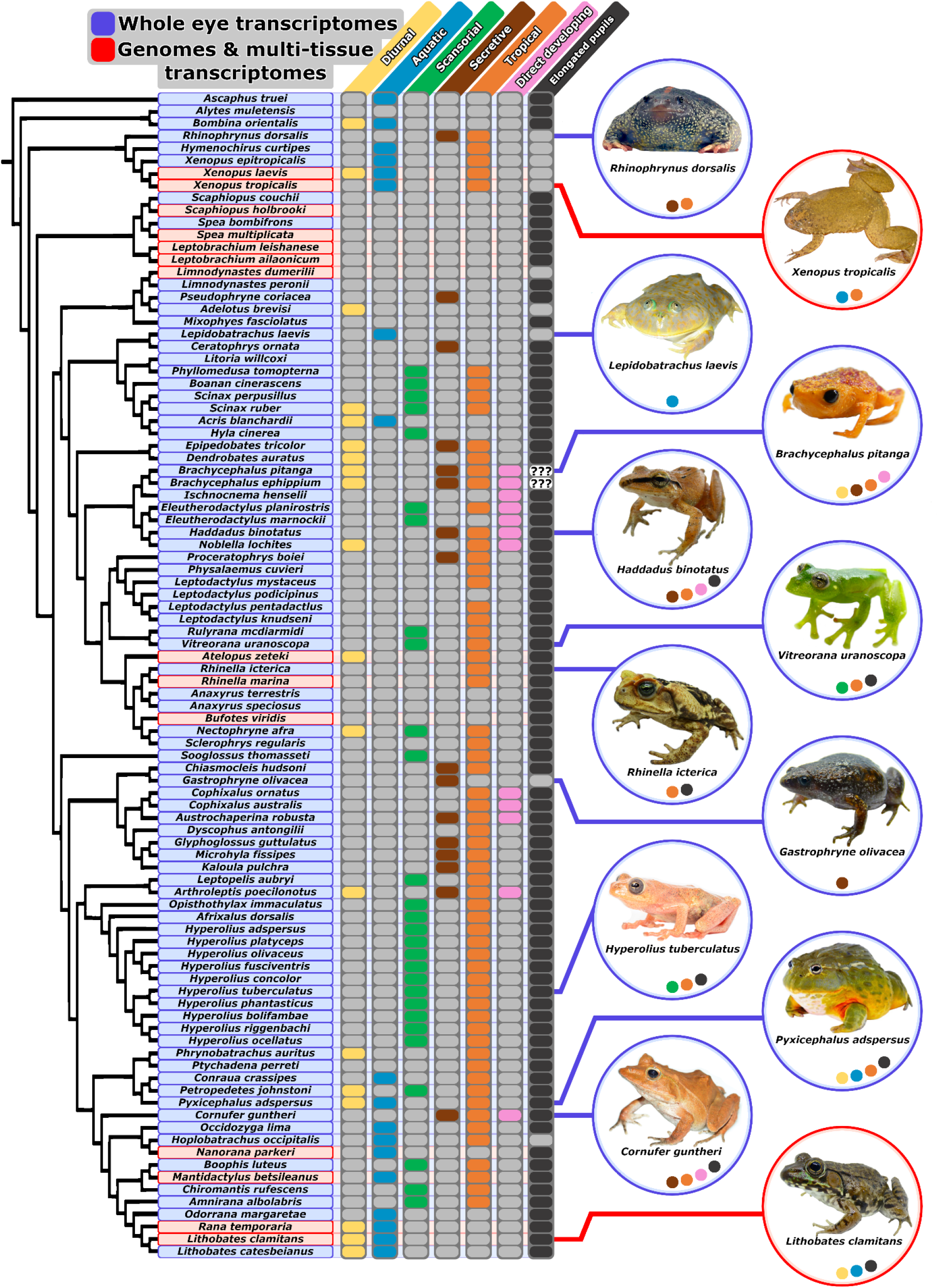
Variation in activity period, habitat, distribution, life history, and pupil shape across our species sampling. Each column represents one of seven trait partitions used to analyze shifts in selective constraint across discrete environmental, developmental, and morphological transitions in frogs. Colored bubbles in trait columns indicate the foreground partition for selection analyses (e.g. diurnal). Gray bubbles indicate the background partition for selection analyses (e.g. non-diurnal). “**???**” indicates that a species was excluded from analysis for a particular trait. Phylogenetic hypothesis based on Feng et al. (2017). *P. adspersus* photo by John Clare

### 2.2 Transcriptome sequencing and assembly

Total RNA was extracted from whole eyes using the Promega Total SV RNA Extraction kit (Promega). Tissue was homogenized in the prepared lysis buffer using the Qiagen Tissuelyzer (10 minutes at 20 Hz). Messenger RNA library construction was performed using the Kapa HyperPrep mRNA Stranded with Riboerase kit (Roche). Each indexed sample was pooled in equimolar amounts and sequenced on two lanes of a NovaSeq4000 with paired end 150 bp reads at the UT Arlington North Texas Genome Center. Prior to assembly, adapters and low-quality bases (q<5) were removed with TrimGalore! (https://www.bioinformatics.babraham.ac.uk/projects/trim_galore/), which implements Cutadapt (Martin 2011). Read pairs shorter than 36bp after trimming were discarded, as were unpaired reads. Quality of processed reads was assessed with FastQC (http://www.bioinformatics.babraham.ac.uk/projects/fastqc/). Transcriptome assembly of each sample was performed *de novo* using Trinity (Grabherr et al., 2011) incorporating all paired reads following the standard protocol.

### 2.3 Non-visual opsin sequence recovery

18 non-visual opsin coding sequences were extracted from frog genomes, consisting of six neuropsins (NEUR1-6), three teleost multiple-tissue opsins (TMT1-3), two melanopsins (OPN4m and OPN4x), encephalopsin (OPN3), parietopsin (PAR), parapinopsin (PARA), vertebrate ancient opsin (VAOP), pinopsin (PIN), retinal g-protein coupled receptor (RGR), and peropsin (RRH), using query sequences from amniotes (Gemmel et al. 2020) and BLAST searches (Altschul, Gish, Miller, Myers, & Lipman, 1990). Complete sequences were compiled and used in turn as queries for BLAST extraction from the frog eye transcriptomes and genomes. Recovered non-visual opsin coding sequences were assembled and aligned against the query sequences in MEGA (Tamura et al., 2011). A coding sequence was considered whole if it was recovered in entirety from start to stop codon, and partial if the sequence was incomplete and more than 50% of codons were recovered. Coding sequences were excluded from analyses if less than 50% of codons were recovered. When present, premature stop codons were converted to sequence gaps to enable their inclusion in downstream analyses (Yohe et al., 2017; Janiak, Chaney, & Tosi, 2018).

### 2.4 Species trait classification

Frog species were partitioned into seven binary trait categories that we predicted to influence the evolution of non-visual light detection. The adult activity period partition included diurnal (including strictly diurnal and mixed diurnal/nocturnal species) vs. non-diurnal species. Our three habitat partitions included aquatic (including semi-aquatic) vs. non-aquatic, scansorial vs. non-scansorial, and secretive (species with lifestyles generally dominated by low-intensity light, including fossorial, burrowing, and leaf-litter species) vs. non-secretive. We used a distribution partition as a proxy for seasonality, including tropical vs. non-tropical species. Our life history partition included direct-developing vs. biphasic species. Finally, our pupil shape partition included species with elongated (horizontal or vertical pupils) vs. non-elongated pupils (other symmetrical shapes). Partitions for all traits are illustrated in Figure 3. We used peer-reviewed literature, online natural-history resources, field guides, and field observations to partition species into these trait categories (Supplementary Table S4). These partitions largely conform to trait scoring in several previous studies of frog visual biology (Thomas et al., 2020, 2021, 2022; Mitra et al., in review), with exceptions noted in Supplementary Table S4. Species that were missing information for a trait were excluded from analyses of that trait.

### 2.5 Selection Analyses

To estimate the strength and form of selection acting on anuran non-visual opsins, each dataset was analyzed using codon-based likelihood models from the codeml program of the PAML 4 software package (Yang, 2007). Coding regions for each of the recovered non-visual opsins were aligned with MUSCLE codon alignment tool in MEGA (Tamura et al., 2011) followed by manual correction. Sequence alignments were imported into BlastPhyMe (Schott, Gow, & Chang, 2019) and maximum likelihood (ML) gene trees were generated using PhyML with default settings (Guindon & Gascuel, 2003). Species topologies were generated through manual editing of PhyML gene topologies to match well-supported phylogenetic relationships among frogs (Feng et al., 2017).

All analyses were performed twice; once using the ML gene-tree topology, and again using the species-tree topology. All topologies were modified to contain the basal trichotomy required by PAML 4 and pruned to match the taxa in a given alignment for each of the 14 non-visual opsins. Trees were then paired with their respective gene alignment and each dataset was analyzed using the PAML 4 random sites models (M0, M1a, M2a, M2a_rel, M3, M7, M8a, and M8), which estimate the rates of nonsynonymous to synonymous nucleotide substitutions (ω, or dN/dS) to infer alignment-wide selection patterns and to test for positive selection acting on non-visual opsin genes. All analyses were run using varying starting values to increase the likelihood of finding global optima. Significance and best fit among model pairs was determined using a likelihood ratio test (LRT) with a χ^2^ distribution.

To test if shifts in selection among non-visual opsins corresponded to variation in adult activity period, habitat, distribution, life history, and pupil shape, we used PAML clade models C and D (CmC and CmD; (Bielawski & Yang, 2004)). These clade models test for evidence of a codon site class demonstrating a shift in selection between pre-partitioned “foreground” and “background” groups (e.g., diurnal frogs and non-diurnal frogs), which can be any combination of branches and clades within a phylogeny. CmC is compared to the null model M2a_rel and assumes that some sites evolve conservatively across the phylogeny (two classes of sites where 0 < ω_0_ < 1 and ω_1_ = 1), whereas a class of sites is free to evolve differently among two or more partitions (e.g. ω_D1_ > 0 and ω_D1_ ≠ ω_D2_ > 0; (Weadick & Chang, 2012)). Alternatively, CmD is compared to the null model M3 and all three site classes (ω_0_, ω_1_, ω_D_) are unconstrained and can assume any value. This can be useful when there is little support for a neutral class of sites. The partition schemes were tested in each non-visual opsin, with each partition corresponding to one of the seven binary trait categories illustrated in Figure 3.

In cases where PAML analyses found significantly elevated ω in a particular group of interest (e.g., elevated ω in diurnal frogs compared to non-diurnal frogs), we used RELAX (Wertheim, Murrell, Smith, Kosakovsky Pond, & Scheffler, 2015), implemented on the Datamonkey web server (Delport, Poon, Frost, & Kosakovsky Pond, 2010), to determine if the elevated ω is the product of relaxed selective constraint (i.e., lack of selection against a change) or adaptive selection (i.e., selection for a change). Such a distinction is useful when attempting to interpret the biological significance of an elevated ω value. RELAX produces a selection intensity parameter, or K value, which modulates the degree to which different ω site classes diverge from neutrality (ω = 1) in pre-partitioned background and foreground groups. When K < 1, this indicates relaxed selection on foreground group branches compared to background group branches. Alternatively, when K > 1, this indicates adaptive selection on foreground group branches compared to background group branches.

## 3 Results

### 3.1 Fourteen non-visual opsins consistently expressed in frog eyes

Across our total sampling, we recovered either whole or partial coding sequences for all 18 non-visual opsins. However, whole or partial coding sequences of NEUR2, OPN3, PAR, and PARA were each recovered in less than 21% of our total samples. OPN3 was not recovered from any of the frog eye transcriptomes, and only two whole coding sequences and four partial coding sequences were recovered across the genomes and multi-tissue transcriptomes (5.61% recovery across our total sampling). Similarly, only one partial coding sequence of NEUR2 was recovered from the frog eye transcriptome sampling, with one whole coding sequence and four partial sequences recovered from the genomes and multi-tissue transcriptomes (5.61% recovery across our total sampling). Coding sequence recovery for PAR and PARA was marginally more successful with 20.56% and 17.76% recovery, respectively, across our total sampling. These four genes (OPN3, NEUR2, PAR, and PARA) were dropped from downstream analyses because their low rates of recovery success limited our ability to generate reliable phylogenies and perform PAML analyses. We recovered the remaining 14 non-visual opsins with some degree of consistency (ranging from 29.9 to 95.3% recovery of whole or partial coding sequences, detailed in Supplementary Table S5) across our total sampling.

### 3.2 Evidence for positive selection in a subset of frog non-visual opsins

To determine overall selective constraint acting on each non-visual opsin, we used the PAML M0 model to average ω across all codon sites in each gene alignment. These tests revealed fairly consistent selective constraint acting on frog non-visual opsins, with most opsins demonstrating mean ω values between 0.09 and 0.19 as illustrated in Figure 4a. Only NEUR6 fell outside of this range, demonstrating an elevated mean ω value of 0.30 (Supplementary Table S6). Taken together, all 14 non-visual opsins have a mean ω < 1, suggesting negative purifying selection. This is expected in most functional protein-coding genes, whose proteins are made up of a high proportion of invariable amino acids (with ω near 0) due to strong functional constraints (Yang, Nielsen, Goldman, & Pedersen, 2000). However, genes demonstrating overall negative selection may still contain positively selected codon sites. We tested for this using the PAML M8 model, which unlike the M0 model, allows ω to vary between sites in a gene. The M8 model is compared with the null model M7 (which allows ω to vary, but constrains ω ≤ 1) to test for the presence of positively selected sites using a likelihood ratio test. Using this approach, we found statistically significant positive selection at a proportion of sites in NEUR1, NEUR6, OPN4x, PIN, and TMT1 as illustrated in Figure 4b. The most extreme signature of positive selection was detected in PIN, which had an ω value of 3.13 (p < 0.001). For comparison, the second most elevated signature of positive selection was observed in NEUR6, with an ω value of 1.67 (p = < 0.001; Supplementary Table S6). These results provide evidence for positive selection in frog non-visual opsins and suggest that adaptive evolution may be occurring within a subset of these genes.

**FIGURE 4.**
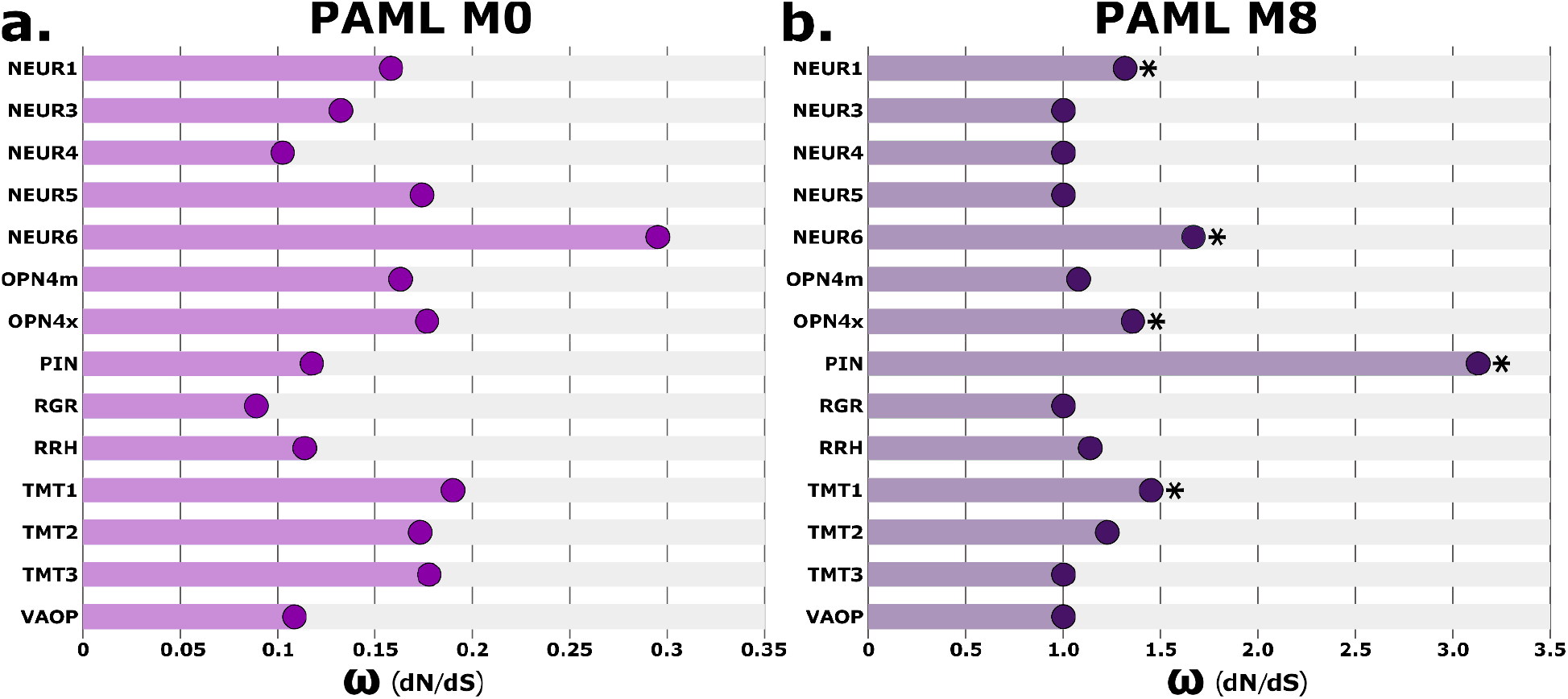
Patterns of selective constraint across non-visual opsin genes. **a.)** The PAML M0 analysis (light purple) averages ω values across all codon sites in a gene alignment. Among non-visual opsins, NEUR6 demonstrated the most elevated ω while RGR demonstrated the least elevated ω, suggesting variation in selective constraint between non-visual opsin genes. **b.)** The PAML M8 analysis (dark purple) tests for the presence of positively selected codon sites in a gene alignment. Five non-visual opsins demonstrated statistically significant (p < 0.05) evidence for positively selected sites (indicated with a “*”). Note that graphs (a.) and (b.) use different scales along the x-axes.

### 3.3 Shifts in selective constraint among non-visual opsins are associated with variation in activity period, ecology, distribution, life history, and pupil shape

Because analyses of gene and species topologies sometimes yielded different significant results, we gave greater weight to results where the same partition was significant across analyses of both topologies and report only those results here. Full results tables are available in the Supplementary Information (Supplementary Tables S7-S10). Significant differences in selective constraint were detected for each of our trait partitions (Figure 5). The diurnal partition was significant in two genes and was the best-fit partition for PIN (ω_diu_ = 0.39 / ω_non-diu_ = 0.26, p = 0.005), indicating that the difference in selective constraint on PIN between diurnal species and non-diurnal species was greater than the difference in any other PIN partition. Among the habitat partitions, the aquatic partition was significant in three genes and was the best-fit partition for NEUR4 (ω_aqt_ = 0.30 / ω_non-aqt_ = 0.23, p = 0.03) and RGR (ω_aqt_ = 0.34 / ω_non-aqt_ = 0.20, p < 0.001). The scansorial partition was significant in two genes and was a best_fit in NEUR5 (ω_scn_ = 0.37 / ω_non-scn_ = 0.26, p = 0.003). The secretive partition was not significant in any genes across analyses of both the gene and species topologies. For the distribution partition, which served as a proxy for differences in seasonality across tropical and non-tropical distributions, we detected significant variation in selective constraint in four genes, including a best fit in NEUR1 (ω_trp_ = 0.26 / ω_non-trp_ = 0.31, p = 0.03). Interestingly, the life-history partition was the most frequent significant partition among non-visual opsins, with six genes demonstrating a significant difference in selective constraint between direct-developing and biphasic species. The direct-developing partition was also the best-fit partition in each of these six genes (NEUR3 (ω_drd_ = 0.43 / ω_lrv_ = 0.27, p = 0.002); NEUR 6 (ω_drd_ = 0.50 / ω_lrv_ = 0.28, p = 0.002); OPN4m (ω_drd_ = 0.50 / ω_lrv_ = 0.29, p < 0.001); RRH (ω_drd_ = 0.38 / ω_lrv_ = 0.25, p = 0.006); TMT1 (ω_drd_ = 0.48 / ω_lrv_ = 0.25, p < 0.001); and TMT3 (ω_drd_ = 0.37 / ω_lrv_ = 0.21, p = 0.03)). Finally, because OPN4m has been implicated in regulating the pupillary light response, we tested the elongated pupil partition to explore how constricted pupil shape might relate more generally to non-visual opsin evolution. Across topologies, the elongated pupil partition was not significant in OPN4m and was otherwise significant in two genes, with no best fits.

**FIGURE 5.**
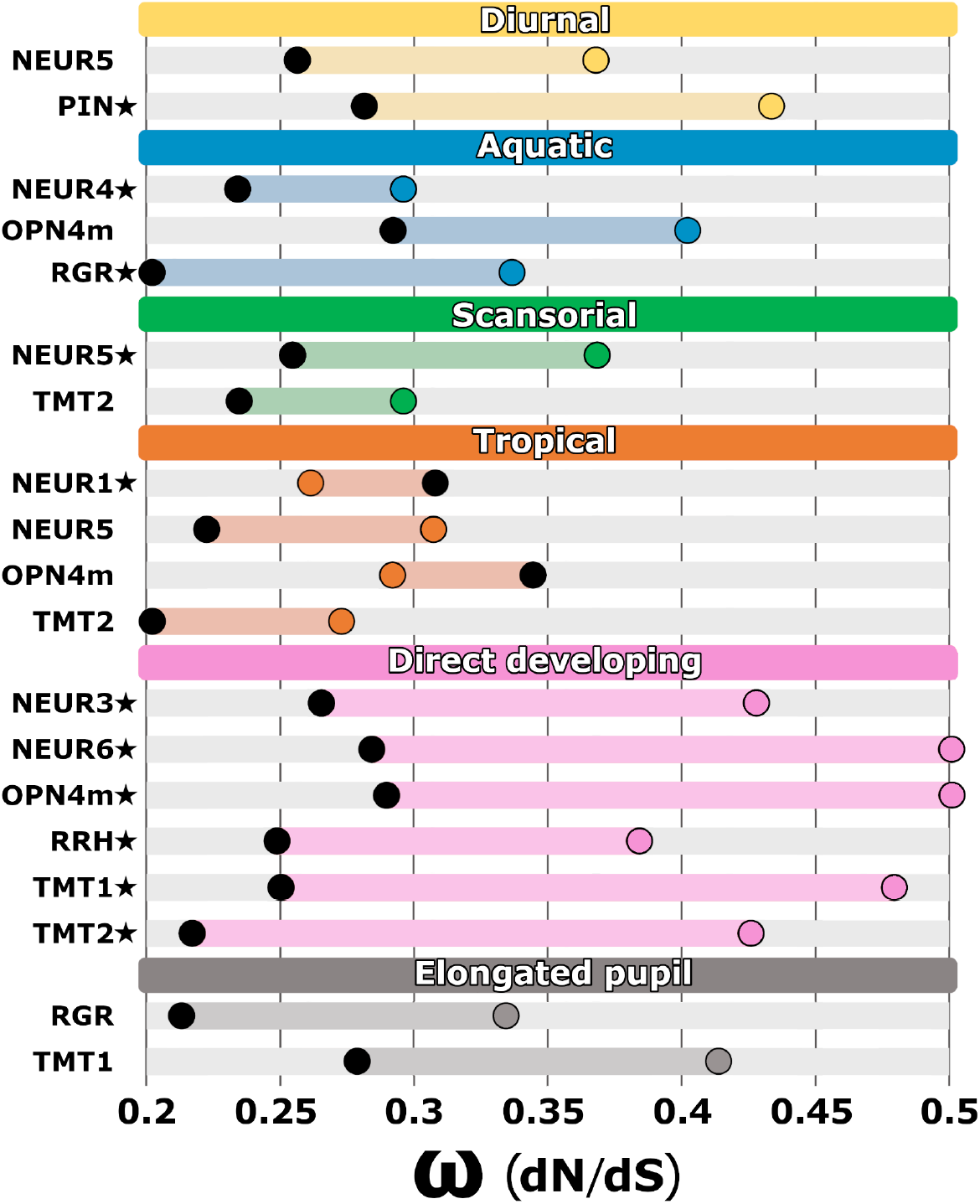
Shifts in selective pressure on non-visual opsin genes across frog trait partitions as illustrated in Figure 2. The ω (dN/dS) values of the divergent site class using CmC analysis of non-visual opsin species topologies are shown, highlighting the difference between the foreground (colored circle) and the background (black circle) partitions for each gene. Under each trait partition, genes with statistically significant (p < 0.05) shifts in selection across analyses of both gene and species topologies are shown. Best fit partitions are indicated with a star. Complete PAML results are available in Supplementary Tables S7, S8, S9, and S10.

We used RELAX to test specific hypotheses (see sections 4.3–4.5) regarding non-visual opsin evolution in genes that have known functions and demonstrated a significant shift in selective constraint across a partition. These analyses revealed evidence for relaxed selection on OPN4m in aquatic (K = 0.47, p < 0.001) and direct-developing species (K = 0.33, p < 0.001), as well as relaxed selection on PIN in diurnal species (K = 0.75, p = 0.002). These analyses also revealed no evidence for relaxed or intensified selection on NEUR1 in non-tropical species (K = 1, p = 0.987). A full summary of our RELAX results is available in Supplementary Table S11.

## 4 Discussion

Using a combination of *de novo* assembled whole-eye transcriptomes and previously published genomic and transcriptomic resources, we obtained non-visual opsin sequences from 98 frog species spanning 35 families. We consistently recovered 14 non-visual opsin genes from frog eye transcriptomes, and positive selection was detected in a subset of these genes, most notably NEUR6 and PIN. We also found variation in selective constraint between frog lineages partitioned by activity period, habitat, distribution, life history, and pupil shape, which may reflect functional adaptation in frog non-visual opsin genes. Below we discuss these findings with respect to our current understanding of non-visual opsin expression and function across diverse taxa.

### 4.1 Unexpected non-visual opsin diversity across frogs

The common ancestor of all vertebrates is estimated to have had a genomic complement of 18 non-visual opsins (Beaudry et al., 2017; Gemmell et al., 2020). However, over the course of vertebrate evolution, non-visual opsin diversity has shifted across groups. Non-visual opsin gene losses have been most apparent in groups with primarily nocturnal evolutionary histories, including between nine and eleven losses in mammals, nine losses in snakes, five losses in geckos, and four losses in crocodilians (Gemmell et al., 2020). These losses have all been attributed to a “nocturnal bottleneck” in which ancestors of each group transitioned to nocturnal lifestyles and encountered a reduced need for broad spectral sensitivity (Borges et al., 2018; Emerling, 2017; Pinto et al., 2019). Adult frogs primarily have nocturnal lifestyles, and this activity period is thought to be the ancestral condition for anurans (Anderson & Wiens, 2017). Given this evolutionary history, we expected frogs to demonstrate reduced non-visual opsin diversity compared to the common ancestor of vertebrates. Instead, we recovered remarkable non-visual opsin diversity across frogs, with 14 of 18 non-visual opsins consistently recovered from eye tissues. Of the four genes we failed to consistently recover, two genes, PAR and PARA, are thought to be expressed exclusively in the pineal region of the brain, and thus it is not surprising that we had limited success recovering these non-visual opsins from our eye transcriptome dataset. The inconsistent recovery of the other two genes, NEUR2 and OPN3, is less straightforward to understand. To our knowledge, there is no published expression profile or functional study of NEUR2 in any taxa, which limits our ability to speculate on why we failed to consistently recover this gene. On the other hand, OPN3 studies in other animals report expression in many tissues, including the retina, brain, and liver (Halford et al., 2001). Thus, the absence of OPN3 in all 88 frog eye transcriptomes is surprising given reports of its expression in eye or retinal tissue across diverse taxa, including human retinas (Halford et al., 2001), chicken retinas (Rios, Marchese, & Guido, 2019), and zebrafish eyes (Davies et al., 2015). Although the retina is reported to have the greatest opsin diversity of any tissue (Davies et al., 2015), some non-visual opsins may be expressed at very low levels (e.g. Do, 2019) and there remains a possibility that both NEUR2 and OPN3 are expressed at levels below our threshold of detection, especially considering that we sequenced whole-eye tissue and not isolated retinal tissue.

Comparatively, across our multi-tissue transcriptome and whole-genome datasets, we had much greater success recovering PAR and PARA (Supplementary Table S5). This suggests that both PAR and PARA are retained in frogs, however these genes appear to be expressed primarily in extra-ocular tissues. We failed to consistently recover NEUR2 and OPN3 across both the multi-tissue transcriptome and whole genome datasets. Across fifteen genomes and four multi-tissue transcriptomes, we recovered one complete sequence of NEUR2 from the *Pyxicephalus adspersus* genome (Denton, Kudra, Malcom, Preez, & Malone, 2018), suggesting that this gene is functional in at least some frog species. We also recovered partial coding sequences (> 50% sequence recovery) of NEUR2 from four frog genomes (Supplementary Table S5). For OPN3, we recovered two complete sequences and four partial coding sequences from the genomes (Supplementary Table S5). Additionally, for both NEUR2 and OPN3, we found no evidence of nonfunctionalization among our recovered sequences. Consequently, we conclude that none of the ancestral vertebrate non-visual opsins have been lost across frogs and instead, frogs appear to have maintained a remarkably diverse repertoire of these genes.

### 4.2 Support for adaptive evolution in non-visual opsins

Frogs have maintained an unexpectedly diverse complement of non-visual opsins, especially for a group demonstrating primarily nocturnal activity patterns. This may be due, at least in part, to the biphasic life history of frogs. Although many fully metamorphosed frogs of biphasic species are nocturnal, this is not necessarily true of their aquatic larvae. Instead, many species that are nocturnal as adults are active and forage in daylight as larvae (Beiswenger, 1977; Ding, Lin, Zhao, Fan, & Wei, 2014; Griffiths & Mylotte, 1988). This dual life history may subject non-visual opsins to disparate environmental constraints and selective pressures across metamorphosis, resulting in adaptive decoupling between diurnal larvae and nocturnal adults (Ebenman, 1992; Schott et al., 2021).

Adaptive decoupling may explain why the diurnal and secretive partitions were only significant for a handful of non-visual opsin genes. These partitions were designed to represent transitions to low-light ecologies across frogs, which we expected to influence non-visual opsin evolution because loss of non-visual opsin diversity is often attributed to low-light adaptation. However, disparate light conditions across metamorphosis may complicate the diurnal and secretive adult partitions, confounding our ability to test expectations of the nocturnal bottleneck hypothesis, which posits low-light adaptation as the most proximate driver of opsin evolution and diversity in many taxa (Borges et al., 2018; Emerling, 2017; Pinto et al., 2019).

Alternatively, our findings may offer support for a different hypothesis of opsin evolution. Beaudry et al. (2017) critiqued the nocturnal bottleneck hypothesis’ focus on evolutionary transitions to low-light ecologies and instead focused on developmental transitions as driving opsin evolution and diversity. They pointed to opsin losses in Mammalia, Aves, and Squamata, emphasizing that these groups undergo much of their development within the dark confines of a womb or shell. By contrast, most amphibians and fishes undergo free-living larval development in relatively bright environments and appear to have retained large opsin repertoires (Davies et al., 2015, this paper). Opsin diversity imparts sensitivity to a broad range of light wavelengths, which is likely beneficial to larval amphibians and fishes, whose translucent or transparent bodies make them particularly susceptible to photo-oxidative stress and DNA damage caused by exposure to light (Beaudry et al., 2017). However, Beaudry et al. (2017) only considered opsin losses in squamata at a coarse taxonomic scale and it should be noted that losses in squamates are restricted to ancestrally nocturnal groups (snakes and geckos; Gemmell et al., 2020). While this should cast doubt on the relevance of egg-based development in driving opsin evolution and diversity, the emphasis on development is still worthwhile. For example, recent annotation of the tuatara genome revealed only one opsin loss despite nocturnal ancestry (Gemmell et al., 2020). This maintenance of opsin diversity was hypothesized to result from the tuatara’s unusual life history, in which juvenile tuatara often adopt diurnal and arboreal lifestyles to avoid predation by cannibalistic adults, which hunt primarily at night. Thus, frogs and tuataras both often experience disparate light environments across development, which may explain why these groups demonstrate the lowest rates of opsin gene loss among amniotes despite nocturnal ancestry.

The importance of development in driving opsin evolution and diversity may explain why the direct-development partition was the most frequently significant partition among non-visual opsins, with six genes demonstrating a significant difference in ω between direct-developing and biphasic frog species. The direct-development partition was also the best fit in these six genes and in each case, the direct-developing species demonstrated elevated ω. This higher rate of nonsynonymous nucleotide substitution in non-visual opsins among direct-developing frogs may be the product of either relaxed or adaptive selection. Considering the Beaudry et al. (2017) hypothesis, we predicted that elevated ω values in direct-developing lineages are driven by relaxed selection on non-visual opsins because these genes are less adaptive in species without larvae. Our RELAX analyses supported this prediction, revealing significant evidence for relaxed selection in all six of the genes with significant direct-development partitions (NEUR3, NEUR6, OPN4m, RRH, TMT1, and TMT2; supplementary table S11). In parallel analyses of frog visual opsins, we did not observe the same pattern of significance across our direct-development partitions (Schott et al., in prep), suggesting that the pattern observed in non-visual opsins is specific to these genes and not necessarily reflective of broadly elevated ω in light-sensing genes across the genomes of direct-developing species. Collectively, these results support the hypothesis that many non-visual opsins are especially relevant in species whose complex life histories expose them to disparate light environments across development.

### 4.3 OPN4, behavioral light responses, and life history

In zebrafish larvae, OPN4 plays a role in triggering non-directional, stochastic hyperactivity in darkness, resulting in the aggregation of larvae into illuminated areas where they remain due to reduced activity (Fernandes et al., 2012). This behavior, known as dark photokinesis, is understood to drive zebrafish larvae out of dark areas, allowing them to maintain a homeostatic distribution in illuminated waters. Similar behavioral light responses have been observed in frog larvae (e.g. Beiswenger, 1977; Branch, 1983; Ding et al., 2014; Fraker, 2008; Jaeger & Hailman, 1976; Muntz, 1963; Roberts, 1978). Given these observations, we hypothesized that OPN4 may play a role in regulating behavioral light responses of frog larvae. We tested this hypothesis using the direct-development partition from our PAML clade-model analyses with the two OPN4 gene families found in frogs. We found a significant difference in selective constraint acting on OPN4m between direct-developing and biphasic frog species, with the former demonstrating elevated ω. Furthermore, our RELAX analysis indicated that this elevated ω is the result of relaxed selection in direct-developing species. Considering our understanding of OPN4’s function in larval zebrafish, the relaxed selection observed in direct-developing frogs may be evidence of reduced functional relevance in species without aquatic larvae. Furthermore, OPN4m is also differentially expressed in leopard frog eyes across metamorphosis, with significantly reduced expression in fully metamorphosed juveniles compared to larvae (Schott et al., 2022). Taken together, these findings suggest that OPN4m function is especially relevant in larval frogs, potentially contributing to behavioral light responses. However, further functional study in a broad diversity of frog species is needed to support this hypothesis.

### 4.4 PIN and low-light photoreception

PIN was first discovered in the chicken pineal gland in 1994, making it the first opsin to be characterized in an extraocular organ (Okano, Yoshizawa, & Fukada, 1994). Because expression of PIN was initially observed only in the pineal gland, the opsin was thought to function strictly in regulating the production and secretion of melatonin (Csernus, Becher, & Mess, 1999). However, low levels of PIN expression were later reported in the outer retina of a gecko (Taniguchi, Hisatomi, Yoshida, & Tokunaga, 2001), and more recently in rod cells in the outer retina of a fish and frog (Sato et al., 2018). In addition to being expressed in rod cells, PIN also exhibits a thermal isomerization rate strikingly similar to that of RH1, the rod visual opsin responsible for high-sensitivity dim-light vision. This observation suggests that, at least in the eyes of fish and frogs, PIN may function as an RH1-like visual pigment contributing to low-light photoreception (Sato et al., 2018). We found a significant difference in selective constraint of PIN between diurnal and non-diurnal frogs, with diurnal species having an elevated ω value. Considering PIN’s hypothesized role in low-light photoreception, we predicted that this elevated ω value is the result of relaxed selective constraint in diurnal species, which are likely less dependent on low-light photoreception to sense their environments. The RELAX analysis supports this hypothesis, revealing significant evidence for relaxed selection in diurnal species and supporting PIN’s hypothesized role in low-light photoreception.

### 4.5 NEUR and seasonality

Tropical species are typically exposed to less seasonal variation in photoperiod, often causing them to rely on humidity cues to synchronize physiological and behavioral changes with seasonality (Canavero & Arim, 2009; Borah et al., 2019). In amphibians, seasonal precipitation has historically received the most attention as a potential cue stimulating physiological and behavioral changes related to reproduction (Duellman & Trueb, 1994; Feder & Burggren, 1992; Stebbins & Cohen, 1997). However, growing evidence indicates that photoperiod sensitivity may be the most proximal factor determining seasonal changes in physiology and activity in amphibians (Canavero & Arim, 2009). Non-visual opsins have been implicated in the seasonal sensitivity of birds (Nakane et al., 2010), and may serve a similar function in frogs. To test if selection signatures support this possibility, the distribution partition scheme was designed to approximate exposure to seasonal variation in tropical versus non-tropical taxa. We found a significant difference in selective constraint of NEUR1 between tropical and non-tropical frogs, with non-tropical species demonstrating elevated ω values.

NEUR1 is reported to regulate seasonal reproductive changes in quail, including thyroid stimulating hormone secretion and subsequent testicular growth (Nakane et al., 2014). As in birds, frogs demonstrate many conspicuous seasonal changes associated with breeding. These changes include hormone secretion and testicular growth (Delgado, Gutiérrez, & Alonso-Bedate, 1989), as well as nuptial pad development (Willaert et al., 2013) and dynamic sexual dichromatism (Bell, Webster, & Whiting, 2017). One particularly well-studied example of seasonal reproductive changes is found in subtropical Leishan mustache toads (*Leptobrachium leishanense*). During the breeding season, males of this species develop mustache-like nuptial spines on their maxillary skin. The development of these spines has been linked to seasonal steroid biosynthesis and thyroid hormone secretion (Li et al., 2019), implicating similar hormonal pathways to those activated by NEUR1 in quail. If NEUR1 is contributing to seasonal sensitivity and reproductive changes in frogs, we might expect that the elevated ω is the product of adaptive selection in non-tropical species due to the greater seasonal variation in photoperiod they experience. However, our RELAX analysis revealed no signature of relaxed or adaptive selection in non-tropical species. This may be because partitioning species as tropical or non-tropical fails to capture exactly which species rely on photoperiod to cue seasonal reproduction. A better test of NEUR1’s relevance to seasonal reproduction might be to partition frogs that are observed to reproduce continuously throughout the year, such as the tropical toad *Duttaphrynus melanostictus* (Jørgensen, Shakuntala, & Vijayakumar, 1986), and frogs that reproduce on a strictly seasonal basis. However, this test would require a more extensive understanding of seasonal reproductive dynamics across frogs than is available in current literature. Given our findings, NEUR1 warrants further study to clarify its role in regulating seasonal reproductive changes and as a candidate for the basis of seasonal sensitivity in frogs.

## 5 Conclusions

Frogs offer a compelling system in which to study the evolution of light sensitivity across diverse ecologies, morphologies, and life-history strategies. We found that frogs have retained a diverse repertoire of non-visual opsins, with 14 genes consistently recovered from frog eye transcriptomes. At the genomic level, frogs appear to have maintained all 18 ancestral vertebrate non-visual opsins and thus demonstrate the lowest documented rate of opsin gene loss among amniote groups. Signatures of positive selection were detected in a subset of these genes. We also found variation in selective constraint between discrete ecological and life-history classes, which may reflect functional adaptation in frog non-visual opsin genes. Our findings also provide genomic support for emerging hypotheses of non-visual opsin evolution, including the role of PIN in low-light photoreception and the adaptive importance of many non-visual opsins in species with complex life histories.

## Acknowledgments

We thank the following field companions who helped obtain specimens for this work: Hannah Augustijnen, Abraham G. Bamba Kaya, C. Guillherme Becker, Gabriela Bittencourt-Silva, Itzue Calviedes Solis, Patrick Campbell, Diego Cisneros-Heredia, Simon Clulow, Mateo Davila, Paul Doughty, Juvencio Eko Mengue, TJ Firneno, Carl Franklin, Philippe Gaucher, Ivan Gomez-Mestre, Shakuntala Devi Gopal, Jon and Krittee Gower, Célio F. B. Haddad, Anthony Herrel, Sunita Janssenswillen, Jim Labisko, Simon Loader, Simon Maddock, Michael Mahony, Renato A. Martins, Matthew McElroy, Christopher Michaels, Nicki Mitchell, Justino Nguema Mituy, Diego Moura, Martin Nsue, Daniel M. Portik, Ivan Prates, Kim Roelants, Corey Roelke, Lauren Scheinberg, Bruno Simões, Ben Tapley, Elie Tobi, Rose Upton, Mark Wilkinson, and Molly Womack. We thank the Gabon Biodiversity Program and Bioko Biodiversity Protection Program for logistical support in the field; Grant Webster, Scott Keogh, Jared Grummer for advice on where to find key species; and Jodi Rowley and Stephen Mahony for exporting tissues for analysis. Sampling was conducted following IACUC protocols (NHMUK, NMNH 2016-012, UNESP Rio Claro CEUA-23/2017, UTA A17.005, ANU A2017/47) and with scientific research authorizations (USA: Texas Parks and Wildlife Division SR-0814-159, North Cascades National Parks NCCO-2018-SCI-0009, Brazil: ICMBio MMA 22511-4, ICMBio SISBIO 30309-12; United Kingdom: NE Licence WML-OR04; French Guiana: RAA:R03-2018-06-12-006; Gabon: CENAREST AR0020/17; Australia: New South Wales National Parks & Wildlife Service SL102014, Queensland Department of National Parks WITK18705517; Equatorial Guinea: INDEFOR-AP 0130/020-2019). This research was supported by grants from the Natural Environment Research Council, UK (NE/R002150/1) and the National Science Foundation (DEB-1655751). J. Boyette was supported by the NMNH Natural History Research Experience REU program (NSF-OCE:1560088).

## Author contributions

RKS, RCB, and JLB conceived and designed the study with input from MKF, KNT, JWS, and DJG. RCB, MKF, JWS, DJG conducted fieldwork. RCB and MKF performed RNA sequencing. JLB and RKS analyzed the data. JLB, RKS, MKF, and RCB wrote the first draft. All authors edited the manuscript. RKS and RCB provided supervision. RCB, DJG, JWS, and MKF provided project administration and acquired the funding. All authors read and approved the final manuscript.

## Data availability statement

The data underlying this article are available in NCBI under Bioproject *tbd* and Zenodo (*doi tbd*), in addition to that available in the article and its additional files. Raw sequence data were deposited in the NCBI Short Read Archive (SRA accession *tbd*).

